# Lysosomal release of amino acids at ER three-way junctions regulates transmembrane and secretory protein mRNA translation

**DOI:** 10.1101/2023.08.01.551382

**Authors:** Heejun Choi, Ya-Cheng Liao, Young J. Yoon, Jonathan Grimm, Luke D. Lavis, Robert H. Singer, Jennifer Lippincott-Schwartz

**Affiliations:** Janelia Research Campus, HHMI, Ashburn VA; Albert Einstein College of Medicine, Bronx NY

## Abstract

One-third of the mammalian proteome is comprised of transmembrane and secretory proteins that are synthesized on endoplasmic reticulum (ER). Here, we investigate the spatial distribution and regulation of mRNAs encoding these membrane and secretory proteins (termed “secretome” mRNAs) through live cell, single molecule tracking to directly monitor the position and translation states of secretome mRNAs on ER and their relationship to other organelles. Notably, translation of secretome mRNAs occurred preferentially near lysosomes on ER marked by the ER junction-associated protein, Lunapark. Knockdown of Lunapark reduced the extent of secretome mRNA translation without affecting translation of other mRNAs. Less secretome mRNA translation also occurred when lysosome function was perturbed by raising lysosomal pH or inhibiting lysosomal proteases. Secretome mRNA translation near lysosomes was enhanced during amino acid deprivation. Addition of the integrated stress response inhibitor, ISRIB, reversed the translation inhibition seen in Lunapark knockdown cells, implying an eIF2 dependency. Altogether, these findings uncover a novel coordination between ER and lysosomes, in which local release of amino acids and other factors from ER-associated lysosomes patterns and regulates translation of mRNAs encoding secretory and membrane proteins.

## Introduction

Endoplasmic Reticulum (ER) is an intricate network of tubules, junctions, and sheets that extends throughout the cytoplasm in virtually all cell types. Studded with ribosomes, ER interacts with a significant proportion of messenger RNAs (mRNAs) in the cell through diverse mechanisms (*1*). A crucial subset of ER-associated mRNAs encodes secretory and integral membrane proteins, constituting about one-third of the human proteome. These mRNAs, referred to here as “secretome” mRNAs, differ from cytosolic protein-encoding mRNAs in that they undergo co-translational translocation. During this process, the nascent signal peptide from the translating ribosome inserts into the Sec translocon on the ER membrane (*2-4*). This positions the secretome mRNA to be physically linked to the ER surface until there is complete termination of translation.

During co-translational translocation, proper coupling of translation and insertion needs to occur for accurate protein folding and maturation processes. Dysregulation of these events, for example by altering elongation rates of nascent peptides, can activate stress response/protein quality control pathways that impede cellular proliferation and growth (*5*). As regulation of translation involves many translation-related factors that often have compartmentalized activities (*6-8*), a relevant question becomes whether translation of secretome mRNAs occurs preferentially on specialized ER subdomains. Classically, the major site of secretome mRNA translation is thought to occur on ER sheets due to their enrichment of ribosomes (*9*). Recent FIB-SEM reconstructions of ER in a variety of cell types, however, has shown significant ribosome abundance on nearly all ER structures, including sheets, tubules, and junctions (i.e., sites where ER tubules intersect) (*10*). These ER-localized ribosomes are likely engaged in the translation not only of secretome mRNAs but also of non-secretome mRNAs (including those encoding cytosolic and mitochondrial proteins) (*11, 12*). It remains unclear, therefore, whether secretome mRNAs translate broadly across the ER surface or only at specific spatial locations.

Here, we investigate this question by examining the spatial distribution and regulation of secretome mRNA translation. Using single molecule imaging to monitor secretome mRNAs and their subcellular location in live cells, we found that translation of secretome mRNAs occurs preferentially on ER membranes adjacent to lysosomes, the major sites of amino acid storage and release in cells. Neutralizing lysosomal pH or inhibiting lysosomal proteases reduced secretome mRNA translation, suggesting that local release of amino acids and other factors from ER-associated lysosomes is important for secretome mRNA translation. The observed increase in secretome mRNA translation near lysosomes was dependent on the ER-resident protein Lunapark, which colocalizes with secretome mRNA translation hotspots on ER. Conversely, knockdown of Lunapark led to decreased translation of secretome mRNAs with no impact on translation of other mRNAs. The effect of Lunapark knockdown on secretome mRNA translation was reversed upon addition of the integrated stress response inhibitor ISRIB. Together, the results reveal an unexpected role of lysosomes positioned near ER junctions that are enriched with Lunapark in the regulation of secretory and transmembrane protein synthesis on ER.

### Visualizing secretome mRNA dynamics on ER membranes

To monitor single mRNAs encoding membrane and secretory proteins on ER (i.e., secretome mRNAs), we utilized MS2 mRNA reporters (*13*) with open-reading frames that encode different transmembrane and luminal proteins fused to GFP (Fig. 1A). As a model secretome mRNA reporter, we used an MS2 mRNA encoding the first 45 amino acids of the Type II membrane protein sialyltransferase (a Golgi enzyme) fused with EGFP (i.e., SiT-EGFP MS2). A Golgi-like distribution of EGFP was indeed seen in cells expressing this reporter, consistent with appropriate translation of the reporter and trafficking of the synthesized protein (Fig. S1A).

**Figure 1.**
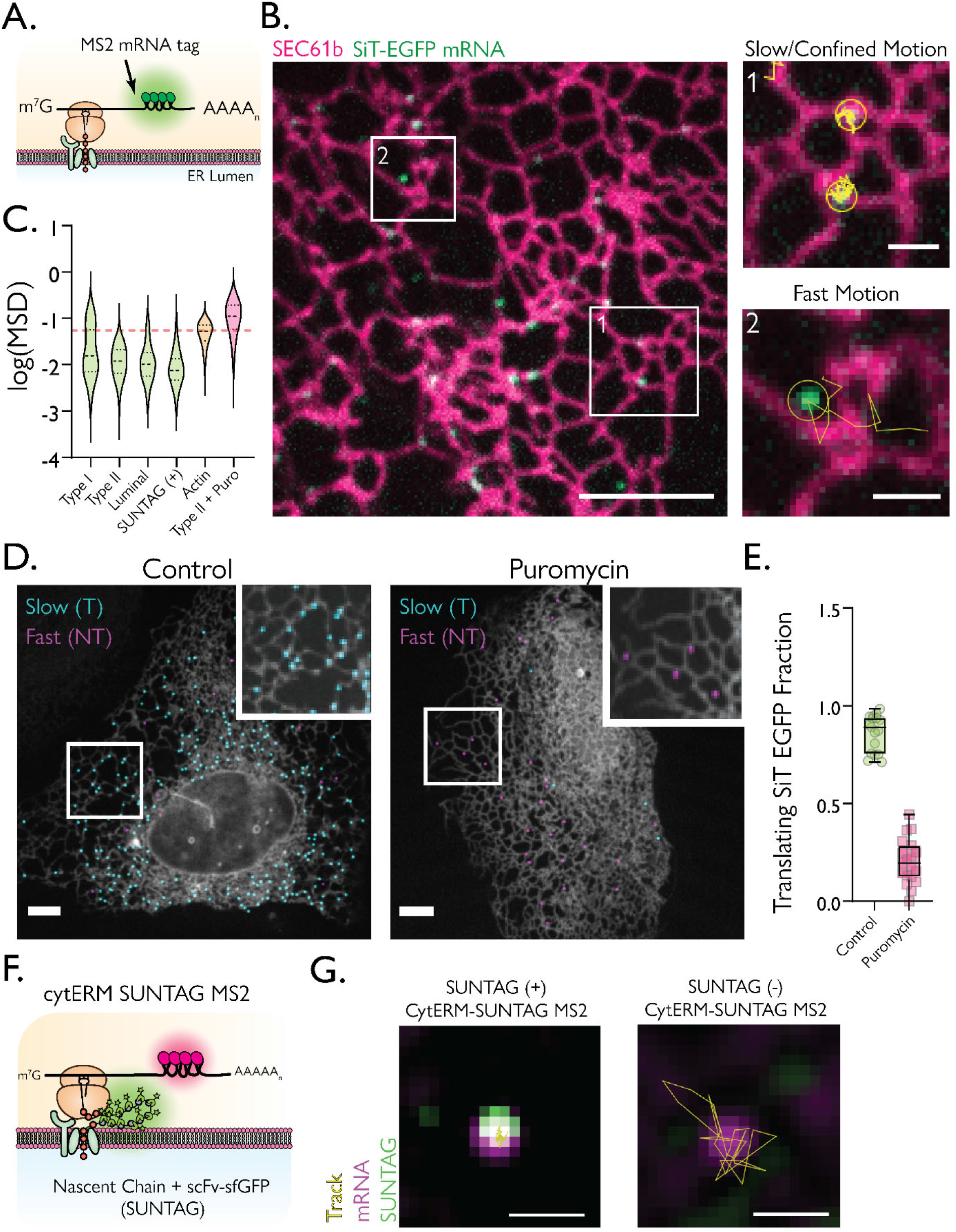
**A)** Design of the mRNA reporter used in the system. The resulting mRNA contains a short 5’UTR, an open reading frame (ORF) of a protein of interest, 24 repeats of the MS2 binding sites (MBS), and a short 3’UTR. The reporter mRNA is transcribed in the nucleus, 5’ capped and 3’ polyadenylated. The transcribed RNA binds MS2 coat proteins (MCP) tagged with a fluorescent protein or the HaloTag. **B)** Spinning disk confocal images of SEC61ß labeling the ER (magenta) and SiT-EGFP mRNA (green). Scale bar = 5 μm. Inset 1 from the overlayed image shows two SiT-EGFP mRNAs that exhibited slow or confined movement as depicted by particle trajectories shown in yellow (Scale bar = 1 μm). Inset 2 shows a single SiT-EGFP mRNA with fast movement near the ER (Scale bar = 1 μm). **C)** Violin plots of log(MSD_t=1sec_), where mean squared displacement (MSD) was in a unit of μm^2^, of (from left to right) Type I (CD4-EGFP), Type II (SiT-EGFP) and Luminal (Calreticulin-mEmerald), SUNTAG(+) (cytERM-SUNTAG MS2), β-actin (Halo-actin), and Type II + puromycin (SiT-EGFP + Puro) mRNAs. The red dotted line indicates the MSD cutoff at 0.055 μm^2^. **D)** Images of control (left) and puromycin-treated (right) U-2 OS cells expressing SNAPf-SEC61ß (gray) and SiT-EGFP mRNAs. Each mRNA is pseudo-colored to represent translationally active (T, blue) or silent (NT, magenta) states. Scale bars = 5 μm. Insets show enlarged views of the cells outlined by white boxes. **E)** Box plots of the fraction of translationally active SiT-EGFP mRNA from control (cyan, N = 17) and puromycin-treated (magenta, N = 19) U-2 OS cells. Each dot represents a cell. **F)** ER-specific translation reporter design. The construct includes rabbit cytochrome p450 signal peptide and transmembrane domains fused with 23 repeats of GCN4 epitope. Single-chain variable fragment (scFv) that recognizes the epitope is tagged with super-folder GFP. It can bind to a nascent chain of GCN4 epitopes and accumulate as a bright punctum (SUNTAG). **G)** Representative merged images of SUNTAG (green), mRNA (magenta), and trajectory (each step = 1 second) (yellow). On the left is a translating SUNTAG(+) particle that exhibits confined motion. On the right is a non-translating SUNTAG(-) particle that exhibits rapid motion. Scale bar = 1 μm.

Visualizing SiT-EGFP MS2 mRNA dynamics in live cells using the ER reporter SEC61ß to label ER membranes, we observed fluorescent puncta representing individual SiT-EGFP mRNAs that were mainly localized on ER membranes (Fig. 1B). Interestingly, the puncta underwent either very slow/confined motion, being stabilized on ER membranes for dozens of seconds (Fig. 1B, inset 1), or displayed rapid movement across the ER signal, coming on/off ER (Fig. 1B, inset 2) (see Movie S1). To test which of these different types of movements correlated with SiT-EGFP MS2 mRNA being actively engaged in co-translational translocation, we treated cells with puromycin, which terminates translation. This dramatically shifted most SiT-EGFP MS2 mRNAs into a fast-moving population with many mRNAs now dislodging from ER (Fig. S1B and Fig. 1C), suggesting that slow-moving SiT-EGFP MS2 mRNA molecules were undergoing translation.

Other secretome mRNAs, including those encoding Type I (CD4-EGFP) and luminal (Calreticulin-mEmerald) proteins, exhibited similar highly confined motion as observed for SiT-EGFP mRNA in untreated cells (Fig. 1C). This contrasted with actin mRNA, which when associated with ER, moved rapidly, exhibiting no confined motion (Fig. S1C and Movie S2). As actin mRNA does not undergo co-translational translocation into ER, the data suggested that the process of co-translational translocation itself restricts the motion of secretome mRNAs.

Given the translocation-dependency of secretome mRNA motion, we used slow/confined motion to identify translating secretome mRNAs. Comparing mRNA motions in control cells with that in puromycin-treated cells, we determined that a mean squared displacement (MSD) of <0.055 μm2 could serve as a cut-off for distinguishing between translating and non-translating mRNAs (Fig. 1C, see red dotted line) (see Fig. S2 for details on the reasoning behind this cutoff). Using this MSD cut-off, we categorized the translation state of SiT-EGFP mRNAs by pseudo-coloring translating mRNA (T) in blue and non-translating mRNA (NT) in magenta based on their MSD (Fig. 1D). Quantification of the data showed that ∼85% of SiT-EGFP mRNA molecules are translating in control cells compared to <15% in puromycin-treated cells (Fig. 1E).

### Only slow-moving secretome mRNAs are engaged in active protein synthesis

To further corroborate the relationship between motion and translation of a secretome mRNA, we used the SUNTAG reporter system, which reads out translation by labeling the nascent peptide (*14-17*). We constructed an ER-specific SUNTAG reporter consisting of the targeting sequence and transmembrane domain of rabbit cytochrome p450 (a secretome protein localized to the ER) fused to GCN4 epitope repeats to visualize nascent peptide, as well as MS2 stem loops to visualize mRNA (i.e., cytERM-SUNTAG MS2) (Fig. 1F). Proper translation of cytERM-SUNTAG MS2 was inferred when fluorescent signal from both mRNA and SUNTAG signals were colocalized.

In cells expressing cytERM-SUNTAG MS2, all fluorescent puncta with colocalized mRNA and SUNTAG signals exhibited highly confined motion (Fig. 1C, and 1G and Movie S3). In contrast, puncta with mRNA fluorescence only (i.e., lacking the SUNTAG signal) moved rapidly across the ER (Fig. 1G and Movie S4). The results demonstrate that secretome mRNAs shift from being highly diffusive when non-translating to being highly confined when translating, thereby confirming that the motion of a secretome mRNA alone can be used to determine whether it is undergoing co-translational translocation.

### ER junctions are hotspots for secretome mRNA translation

We next investigated whether secretome mRNAs are preferentially translated at specific ER subdomains. The ER is a complex membrane system comprised of sheets and tubules that intersect at varying densities ranging from discrete three-way junctions to dense groups of junctions (matrices) to fused junctions forming sheet-like structures (*18*). To test which of these structures serve as ER subdomains for secretome mRNA translation, we imaged secretome mRNAs in the cell periphery where we could optically resolve specific ER morphologies. Translating secretome mRNAs were identified as mRNAs on the ER that had very low mobility (i.e., MSD <0.055 μm^2^ or undergoing little motion for tens of seconds, see above). Interestingly, the translating secretome mRNAs were found primarily at ER junctions. This was seen for secretome mRNAs encoding three different types of proteins, including the Type I membrane protein CD4 (Fig. 2A CD4-EGFP MS2), the Type 2 membrane protein SiT (Fig. 2A SiT-EGFP MS2) and the luminal protein Calreticulin (Fig. 2A, Calreticulin-mEmerald MS2). The translating fraction (T) of these secretome mRNAs were all predominantly localized within 0.2 μm of an ER junction (Fig. 2C, Type I (T), Type II (T) and Luminal (T)). By contrast, non-translating secretome mRNAs (NT), identified by their significantly higher mobility on ER, showed no enrichment at ER junctions, as quantified for the mRNA encoding SiT-EGFP (Fig. 2C SiT (NT)). The average distance of non-translating SiT-EGFP MS2 mRNA from an ER junction resembled that for Halo-actin mRNA on ER, which does not undergo co-translational translocation (Fig. 2C, Halo-actin). The preferred localization of translating secretome mRNAs to ER junctions was independent of cell-type, with slow-moving pools of SiT-EGFP mRNA seen at ER junctions in U-2 OS, HeLa, COS-7, and HT1080 (Fig. S3).

**Figure 2.**
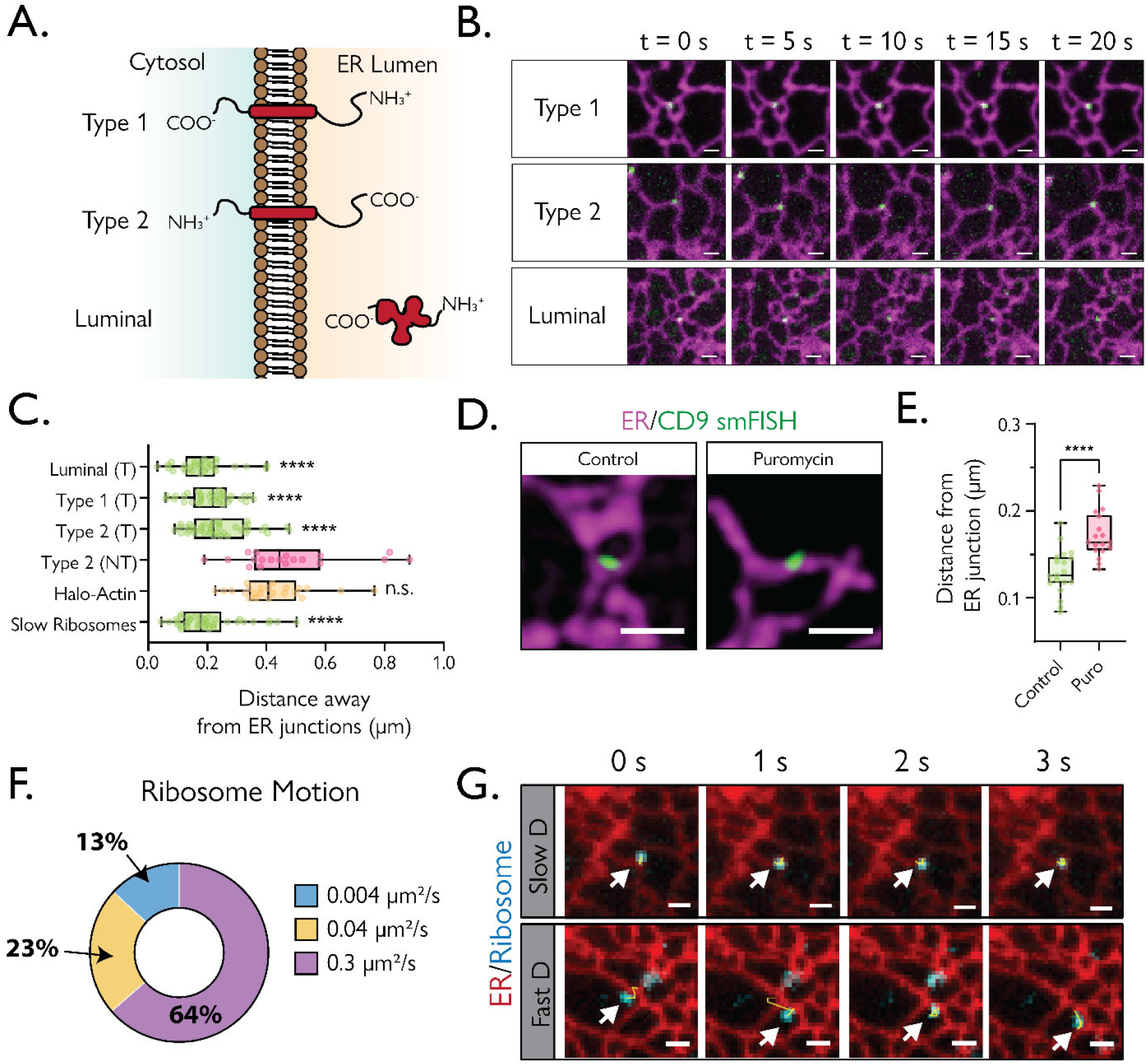
**A)** Topology of Type I, Type II, and ER luminal proteins on the ER membrane. The ER lumen is shown in beige and the cytosol in green. **B)** Time-lapse Images of translationally active mRNAs of Type 1 (CD4-EGFP), Type 2 (SiT-EGFP), and Luminal (Calreticulin-mEmerald) protein mRNAs (green) on the ER (SEC61ß, magenta) in U-2 OS cells. Scale bar = 1 μm. **C)** Box plots of the average distance to the nearest ER junction per individual cell (μm) for translating (T) Luminal (Calreticulin-mEmerald, N = 29), translating Type 1 (CD4-EGFP, N = 32), translating Type 2 (SiT-EGFP (N = 40), non-translating (NT) Type 2 (SiT-EGFP), Halo-Actin mRNA (N = 23), and slow ribosomes (D_eff_ <0.01 μm^2^/s from top to bottom, N = 37). Statistical significance was derived by comparing each <r> distribution with Type 2 (NT). **D)** Representative 3D Structure Illumination Microscopy (3D-SIM) images of hybridization chain reaction single-molecule fluorescent in situ hybridization (HCR smFISH) labeling of endogenous CD9 mRNA (green) overlaid on ER (mEmerald-SEC61ß) in control (left) and puromycin-treated (right) U-2 OS cells. Scale bar = 1 μm. **E)** Quantification of the distance from the nearest ER junction to smFISH spot per individual cell in control (black, N = 19) or puromycin-treated conditions (magenta, N = 18). Each dot represents an average distance from a cell **F)** Circular graph representing the population of diffusion coefficients based on cumulative distribution function fitting of single-step displacement of individual ribosomes. **G)** A montage of a single ribosome (cyan) that has an apparent diffusion coefficient of <0.01 μm^2^/s (Slow D, top) and >0.01 μm^2^/s (Fast D, bottom). Scale bar = 1 μm.

We next examined whether the sites of endogenous secretome mRNA translation were also preferentially localized at ER junctions. CD9 mRNA was used as a representative endogenous secretome mRNA as it is highly expressed in most cell types (*19*). To visualize endogenous CD9 mRNA, we performed hybridization chain reaction single-molecule fluorescent *in situ* hybridization (HCR smFISH) on CD9 mRNAs in U-2 OS cells expressing mEmerald-SEC61ß as an ER marker. Because HCR smFISH does not distinguish translating and non-translating mRNAs, we compared their localization to nearest ER junctions with or without puromycin treatment using 3D structural illumination microscopy. The results showed a clear difference: when translation was inhibited by puromycin, the distance between endogenous CD9 mRNAs and ER junctions increased (Fig. 2D, E). This suggested that endogenous CD9 mRNA translation preferentially occurs at ER junctions.

To further support the notion that ER junctions serve as hotspots for secretome mRNA translation, we performed single-molecule tracking of ribosomes by tracking the large ribosomal subunit protein L10A tagged with HaloTag in cells co-labeled with an ER marker. We reasoned that if translation of secretome mRNAs were occurring at ER junctions, then there should be populations of ribosomes at ER junctions exhibiting the same slow motion as the secretome mRNA. Cumulative distribution function fitting revealed three distinct diffusive populations of ribosomes on ER, including fast-, medium- and slow-moving pools (Fig. 2F), but only the slow-moving ribosome pool was localized preferentially to ER junctions (Fig. 2G) and its effective diffusion coefficient (i.e., ∼0.004 μm^2^·s^-1^) matched well with diffusion coefficient measurements for translating secretome mRNAs (see Fig. S1D, *D*_effective_ = 0.007 μm^2^·s^-1^). Thus, slow-moving pools of both translating secretome mRNAs and ribosomes are localized at ER junctions.

### Lunapark marks ER junctions engaged in secretome mRNA translation

ER junctions are thought to form through the fusion of a tip of an ER tubule with another ER tubule whose structure is stabilized by an ER-resident transmembrane protein called Lunapark (*20, 21*). Lunapark is present on a subset of ER junctions (*20*), which we verified through expression of Lunapark-GFP (Fig. 3A). To test whether the ER junctions enriched in Lunapark are preferentially engaged in secretome mRNA translation, we expressed the ER translation reporter (cytERM-SUNTAG MS2) and immunostained for endogenously expressed Lunapark. Notably, only the translating fraction of cytERM-SUNTAG MS2 that were positive for both mRNA and SUNTAG-bound nascent peptide colocalized with Lunapark signal. The control reporter SUNTAG(-) cytERM mRNA puncta exhibited little or no colocalization with Lunapark (Fig. 3B). Using SUNTAG signal to define translating secretome mRNAs, we found a significantly greater amount of translating (T) CytERM-SUNTAG MS2 mRNAs within 300 nm of Lunapark signal compared to non-translating (NT) mRNA molecules (Fig. 3C). Together, the enrichment of translating secretome mRNAs at Lunapark-containing ER junctions suggests that Lunapark may play an important role in secretome mRNA translation.

**Figure 3.**
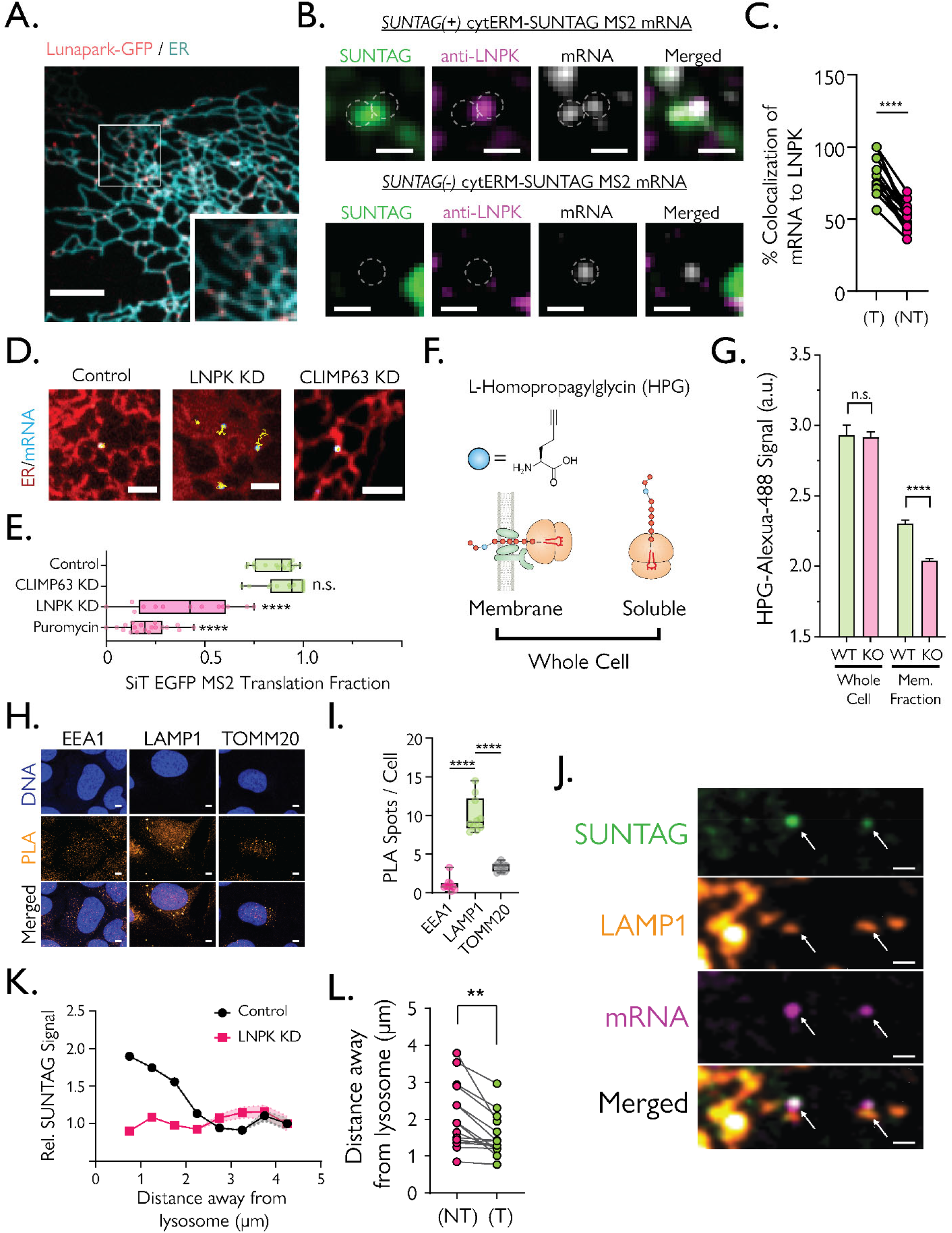
**A)** Representative image of Lunapark-GFP (red) and ER (Halo-SEC61ß, cyan). Inset depicts an enlarged view of the area outlined in a white box. Scale bar = 5 μm. **B)** Top: Representative image of translating SUNTAG(+) signal that overlaps with LNPK (magenta) and cytERM-SUNTAG MS2 mRNA (gray). Bottom: The non-translating cytERM-SUNTAG MS2 mRNA (gray) does not overlap with SUNTAG(-) or LNPK. Scale bar = 1 μm. **C)** Quantification of co-localization of mRNA within a 300 nm radius of segmented Lunapark puncta that are SUNTAG (+) (T) and SUNTAG(-) (NT). Each line represents the same cell. N = 14 **D)** Representative images of SiT-EGFP mRNA (cyan) and particle trajectory (yellow) on the ER (red) from control, Lunapark knockdown (LNPK KD), and CLIMP63 knockdown (CLIMP63 KD) cells. Scale bar = 1 μm. **E)** Translation fraction of SiT-EGFP mRNAs in control (N = 17), CLIMP63 KD (N =14), LNPK KD (N = 13), and puromycin (N = 17) conditions (top to bottom). Statistical comparison was performed against the control distribution. **F)** Depiction of L-homopropargylglycine (HPG) incorporation assay. Whole-cell fraction represents the combined levels of HPG incorporation from membrane and soluble fractions. **G)** Click-labeled Alexa-488-HPG signal from the whole-cell fraction and digitonin-extracted membrane fraction from WT (green) and Lunapark KO U-2 OS (magenta) cells. **H)** Representative images from proximity ligation assay (PLA). DNA (blue), PLA spots (orange), and merged images from PLA reactions with LNPK antibodies against anti-EEA1 (left, early endosome), anti-Lamp1 (late endosome/lysosome), and anti-TOMM20 (mitochondria). Scale bar = 5 μm. **I)** Quantification of PLA spots per cell from anti-EEA1 (left), anti-LAMP1 (middle), and anti-TOMM20 (right) acquired from 10 field of views from 3 replicates. **J)** Representative images of SUNTAG (green), LAMP1 (orange), cytERM-SUNTAG MS2 mRNA (magenta), and merged images. Arrows indicate the location of active translating mRNAs. Scale bar = 1 μm. **K)** Relative SUNTAG intensity binned by the average distance away from the nearest lysosome (μm) in increments of 500 nm for cytERM-SUNTAG MS2 mRNA from control (black) and Lunapark KD (magenta). The shaded area represents the 95% confidence interval. **L)** The average distance between SiT-EGFP mRNA and the nearest lysosome (μm) of translating (T, green) and non-translating (NT, red) mRNA. Each line represents the same cell. N = 15 cells

### Lunapark knockdown selectively reduces secretome mRNA translation

To test whether the presence of Lunapark is necessary for secretome mRNA translation at ER junctions, we knocked down Lunapark (LNPK-KD) in cells expressing SiT-EGFP mRNA. SiT-EGFP mRNA molecules in LNPK-KD cells exhibited a marked increase in overall motility (Fig. 3D). Applying the MSD cut-off value of <0.055 μm^2^ used earlier for identifying translating secretome mRNAs, we found that the pool of translating SiT-EGFP mRNA shifted from 80% in control cells to 40% in LNPK-KD cells (Fig. 3E). A similar result was seen for cytERM-SUNTAG MS2 mRNA using SUNTAG signal as a translation readout (Fig. S4). By contrast, knockdown of the ER-shaping protein, CLIMP63 (*9*), did not alter the distribution/motion of SiT-EGFP mRNAs on ER, with >80% of these mRNAs still undergoing translation (Fig. 3E and S4).

We next tested the impact of Lunapark knockout (LNPK-KO) on global secretion using an imaging method involving incorporation of the amino acid analog L-homopropargylglycine (HPG) into newly synthesized proteins, as analogous to [^35^S]methionine (Fig. 3F) (*22*). Imaging the HPG signal in intact cells, we found no detectable difference in total HPG incorporation in WT versus LNPK-KO cells measured by the clicked Alexa 488 signal (Fig. 3G, left). However, upon digitonin permeabilization of cells to remove the cytosolic fraction, we found a significant decrease in HPG incorporation into proteins retained in membranes and organelle lumens in LNPK-KO cells compared to WT cells (Fig. 3G, right). This suggested the absence of Lunapark lowers the efficiency of membrane and secretory protein synthesis in cells.

### Lysosomes facilitate secretome mRNA translation at Lunapark-enriched ER junctions

Prior work has suggested that lysosomes can be found near sites of translation in axons (*23*), and that lysosomes can associate with ER junctions (*24, 25*). Given these observations and our finding that Lunapark knockdown decreases secretome mRNA translation, we wondered whether lysosome proximity could facilitate secretome mRNA translation at Lunapark-enriched ER junctions. To address this possibility, we first examined whether lysosomes are near Lunapark puncta on ER by performing proximity ligation assays (PLAs) using an anti-Lunapark antibody against antibodies recognizing different organelles, including early endosomes (anti-EEA1), late endosomes/lysosomes (anti-LAMP1), or mitochondria (anti-TOMM20). Significantly higher levels of PLA puncta were detected using LAMP1 antibodies relative to EEA1 or TOMM20 (Fig. 3H and 3I), suggesting lysosomes are near regions of ER junctions enriched with Lunapark.

We next asked whether lysosomes are present near sites of active secretome mRNA translation. For this, we examined the distribution of cytERM-SUNTAG MS2 and fluorescently-labeled LAMP1 in cells. Strikingly, translating cytERM-SUNTAG MS2 puncta (with both SUNTAG and mRNA signals) could be seen juxtaposed with LAMP1-positive structures (Fig. 3J), with the SUNTAG signal from cytERM-SUNTAG MS2 becoming brighter when the reporter was closer to lysosomes (Fig. 3K). This suggested that lysosomes mark hotspots of increased secretome mRNA translation on ER. Supporting this, we found that SiT-EGFP-MS2 mRNA showed highly-confined translating pools (T) significantly closer to lysosomes than that of rapidly-moving non-translating pools (NT) (Fig. 3L).

We then assessed the role of Lunapark in facilitating the distance dependence of secretome mRNAs to lysosomes for efficient secretome mRNA translation. When Lunapark was knocked down, this distance dependency to lysosomes was lost, as shown with our cytERM-SUNTAG MS2 reporter, where the level SUNTAG signal from translating mRNA molecules was now lower and independent of lysosome location (Fig. 3K). Altogether, the above results suggest that secretome mRNA translation is significantly enhanced when lysosomes are near ER junctions enriched with Lunapark.

### Boosted secretome mRNA translation near lysosomes requires lysosomal release of amino acids

Lysosomes are acidic compartments that function to degrade engulfed proteins into amino acids which are then released into the cytoplasm (*26*). Given this, we asked whether lysosomes could serve as local sources of amino acids to facilitate secretome mRNA translation. To test this, we treated cells with inhibitors of lysosomal proteases, or raised lysosomal pH using chloroquine to lower the efficiency of protein degradation in lysosomes. When we examined the amount of translating SiT-EGFP mRNAs under either of these conditions, we found the level was significantly reduced (Fig. 4A). This supported the idea that lysosomal degradation of proteins helps optimize secretome mRNA translation on ER.

**Figure 4.**
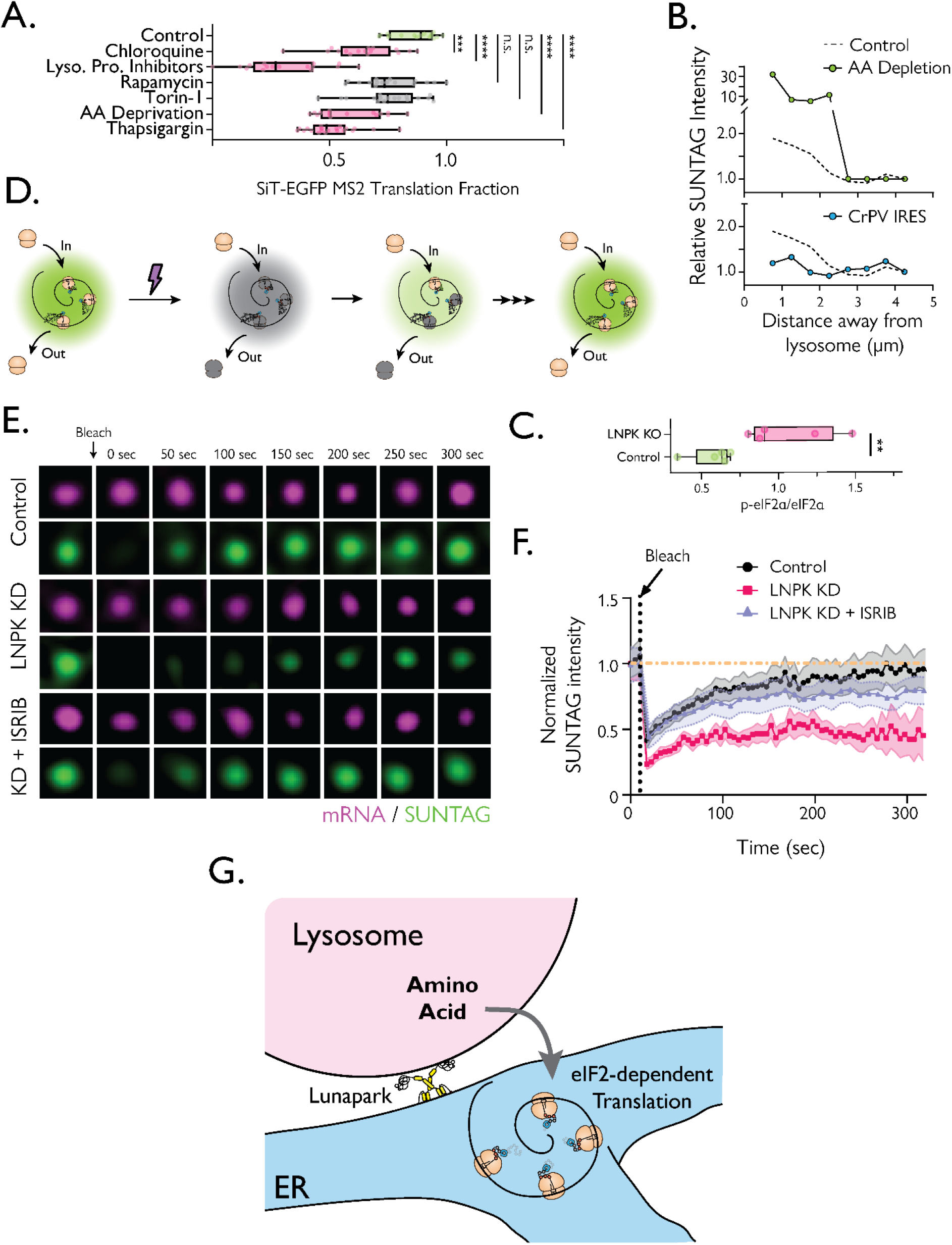
**A)** Boxplot of translating fraction of SiT-EGFP mRNAs under control, chloroquine (10 μM, 4 hr, N = 16), lysosomal protease inhibitors (1 hr, N = 16), rapamycin (100 nM, 1 hr, N = 16), Torin-1 (1 μM, 1 hr, N = 17), amino acid starvation (16 hr, N = 15), and thapsigargin (1 μM, 1 hr, N = 20) treatments. Each statistical comparison was performed against the control. **B)** Relative SUNTAG intensity binned by the average distance away from the nearest lysosome in increments of 500 nm for cytERM-SUNTAG MS2 mRNA under amino acid depletion (AA Depletion, green dot, top) or 5’UTR containing cricket paralysis virus IRES (CrPV IRES, blue dot, bottom) conditions plotted alongside control as shown as a black dotted line. **C)** Boxplot of phospho-eIF2α to eIF2α ratio quantified from immunofluorescence signals from control and Lunapark KO (LNPK KO) U-2 OS cells from 5 replicates. **D)** Schematics of FRAP assay of SUNTAG foci. **E)** Montage of fluorescence recovery of SUNTAG (green) signal from cytERM-SUNTAG MS2 mRNA (magenta) in control (top) and Lunapark knockdown (LNPK KD, middle) and Lunapark knockdown + 200 nM ISRIB (KD + ISRIB, bottom). The arrow indicates photobleaching. **F)** Plot of FRAP recovery curves from control (black), Lunapark Knockdown (LNPK KD, red), and Lunapark Knockdown + 200 nM ISRIB (LNPK KD + ISRIB, purple). The vertical dotted line represents time at photobleaching. The yellow horizontal dotted line represents normalized intensity value at 1. The shaded areas represent 95% confidence intervals for each plot. **G)** Hypothetical model depicting Lunapark-lysosome interaction at an ER three-way junction. The amino acids released by the lysosome can modulate the phosphorylation of eIF2α to promote efficient translation of secretome mRNA.

Lysosomal degradative activity becomes particularly important under amino acid starvation conditions, as the cell now becomes dependent on amino acids from lysosomal degradative activity for protein synthesis (*27, 28*). Examining the amount of translating SiT-EGFP mRNAs under amino acid starvation, we found the overall level was reduced compared to the control (Fig. 4A).

To assess whether secretome mRNA translation becomes more sensitive to lysosome proximity when cells are deprived of amino acids, we used the cytERM-SUNTAG MS2 system to examine whether SUNTAG signals became more closely associated with lysosomes under amino acid deprivation. We found that the distance dependency of secretome mRNA translation to lysosomes under amino acid deprivation was tighter and that the relative SUNTAG signal from cytERM-SUNTAG MS2 was significantly higher in mRNA puncta nearby lysosomes (Fig. 4B). These findings suggested that when amino acids become scarce, secretome mRNA translation now becomes more dependent on nearby lysosomes.

### Secretome mRNA translation responds to the integrated stress response

Amino acid starvation triggers the integrated stress response (ISR), which reduces translation initiation through phosphorylation of eIF2α, a translation initiation factor (*29*). To test whether ISR activation can modulate secretome mRNA translation initiation, we treated cells with thapsigargin, a known activator of the ISR, and examined its effect on SiT-EGFP MS2 translation. Not only did thapsigargin treatment reduce the translating fraction of SiT-EGFP MS2 (Fig. 4A), but this was reversed upon addition of the ISR inhibitor ISRIB (*2*) (Movie S5 and S6). These results suggest that activating the ISR can reduce secretome mRNA translation.

To further examine this phenomenon we modified the 5’ untranslated region (UTR) of cytERM-SUNTAG MS2 to include the cricket paralysis virus internal ribosome entry sequence (CrPV IRES), which allows escape from virtually all regulation of translation initiation, including eIF2-dependent pathways (*30, 31*). When we expressed this construct in cells, we found much lower SUNTAG signal near lysosomes (Fig. S5) and that the relative SUNTAG signal was no longer brighter as they became closer to the lysosomes (Fig. 4A, bottom). It is likely that translation initiation of secretome mRNA translation is regulated by an eIF2-dependent mechanism. In contrast to eIF2 involvement in regulating secretome mRNA translation, a role for mTOR activity was not found. Indeed, inhibiting mTOR activity using rapamycin or Torin-1 treatment had no significant effect on the translating fraction of SiT-EGFP MS2 mRNAs (Fig. 4A). This finding supports prior work showing secretome mRNAs lack the mTOR-responsive 5’ UTR TOP-element that reduces ribosome loading density on mRNA during mTOR inhibition (*32*).

### Dissecting Lunapark’s role in eIF2-dependent secretome mRNA translation

We next examined whether Lunapark played any role in modulating secretome mRNA translation at the level of eIF2 regulation. Phosphorylation of eIF2α is the putative mechanism to repress translation during ISR. When we examined the ratio of phosporylated-eIF2α to eIF2α in control and LNPK-KO cells by immunoblotting and immunofluorescence, we found that while total eIF2α levels were decreased in Lunapark-knockout cells (Fig. S6), the relative amount of phosphorylated eIF2α had increased (Fig. 4C and S6). These findings show that depletion of Lunapark can lead to greater phosphorylation of eIF2α.

To further study this mechanism, we examined the rate of translation through fluorescence recovery after photobleaching (FRAP) experiments in cells expressing the cytERM-SUNTAG MS2 reporter. Individual puncta with mRNA and SUNTAG signal were identified and the SUNTAG was selectively photobleached and monitored for recovery (Fig. 4D). In control cells, photobleaching of SUNTAG puncta led to recovery half-time of 100 seconds (Fig. 4E and 4G black). By contrast, when we photobleached cells with Lunapark knocked out, the cytERM-SUNTAG signal recovered more slowly and never reached pre-bleach levels over 300 seconds (Fig. 4E and 4G magenta). This indicated that the presence of Lunapark was critical for maintaining proper rates of translation of secretome mRNA.

To test if the reduced rates of translation seen in Lunapark knockdown cells is due to phosphorylation of eIF2α, we repeated the experiment but now added ISRIB, which prevents phosphorylated eIF2α from blocking eIF2B GEF function through bypassing the translation repression effect of phosphorylated eIF2α (*33*). Notably, our FRAP experiments examining recovery of the SUNTAG signal after photobleaching showed that ISRIB-treatment increased the recovery rate of the SUNTAG signal to that observed in control (Fig. 4E and 4G Purple). Together, these results strongly support the idea that the effect of Lunapark knockdown on reducing secretome mRNA translation occurs through the regulation of eIF2.

## Discussion

Where mRNA translation occurs within the cell and how it is regulated are longstanding questions necessary for understanding the spatial control of gene expression. In the case of mRNAs encoding membrane and secreted proteins (i.e., secretome mRNAs), the mRNA needs to be directed and targeted to sites on the ER where the nascent peptides insert through the membrane to reach the ER lumen for folding and maturation. To monitor this process in living cells, we visualized mRNAs and nascent peptides simultaneously to localize the site(s) of translation of secretome mRNAs on peripheral ER. Our results revealed a surprising mechanism for regulation of co-translational translocation involving coordination among ER junctions, Lunapark, and lysosomes.

Prior to initiating co-translational translocation of peptide, we found secretome mRNAs diffused freely across ER membranes. Once co-translational translocation began, the mRNAs slowed their motion and became localized to specific ER subdomains known as ER junctions. As ER junctions have less curvature and a relatively larger surface area compared to surrounding ER tubules (*34*), they are ideal sites for formation of large polysome arrays. Similar previous reasoning has led to the belief that stacked ER sheets with ribosomes in the perinuclear region are sites of membrane protein synthesis. Our data showing that ER junctions in the cell periphery are hotspots for secretome mRNA translation indicates that ER sheets are not the exclusive site of synthesis of transmembrane and secretory proteins. In fact, we found that the spacer protein for ER sheets, CLIMP63, was not required for secretome mRNA translation whereas the junction-stabilizing protein, Lunapark, was pivotal for translation regulation.

Lunapark is a zinc-finger containing ER transmembrane protein that localizes to and helps stabilize ER junctions (*20, 21*). Other known roles of Lunapark include it playing a role in ER-phagy (*35*) and in sequestering KLHL12, an adaptor protein for the CUL3 E3 ligase complex (*36*). We found that it was ER junctions marked by Lunapark that were predominantly engaged in secretome mRNA translation and that when Lunapark was knocked out, overall levels of translation of secretome mRNA decreased. Lunapark’s role in regulating translation of secretome mRNAs appeared to be related to eIF2 function. Supporting this, we observed increased phosphorylation of eIF2α (which attenuates translation and triggers ISR) when Lunapark is knocked out. We also saw that reduced secretome mRNA translation in Lunapark knock down cells was reversed by the ISR inhibitor ISRIB.

Proximity ligation assay revealed that lysosomes reside near Lunapark-enriched junctions of the ER, suggesting that secretome mRNA translation may be coordinated with lysosomes. When lysosomes were adjacent to ER junctions, they marked sites of increased secretome mRNA translation. Indeed, ribosome density on secretome mRNAs at ER junctions appeared to increase the closer the mRNA was to lysosomes. This effect was more pronounced during amino acid starvation and was diminished with Lunapark depletion or upon bypassing translation initiation regulation using the CrPV IRES-containing reporter.

What role do lysosomes at Lunapark-enriched ER junctions have in facilitating translation? We hypothesize that secretome mRNAs are directly influenced by lysosomal nutrient sensing through interactions between ER junctions and lysosomes. This interaction could activate known translation-related factors on lysosomes, such as tRNA synthetases (*7*), and other initiation factors (*37*) to modulate translation locally. Supporting this thinking, when we inhibited lysosomal proteases or raised lysosomal pH in cells to block lysosomal degradation enzymes, secretome mRNA translation was significantly reduced. Moreover, the distance dependency of secretome mRNA translation to lysosomes became tighter in cells that were starved of amino acids.

Exactly how lysosomes become associated with Lunapark-enriched ER junctions is unclear. Lunapark could directly associate with a specific lysosomal membrane protein or do so indirectly. Either way, the net result would be lysosomes coming into the vicinity of ER junctions so their release of amino acids and other proteins involved in translation could be used to help boost translation on ER in an eIF2-dependent manner. The presence of Lunapark and lysosomes at ER junctions may be particularly important for translation in regions of the cell far away from the nucleus, where precise spatial control of gene expression becomes critical. Indeed, in neurons, lysosomes ferry RNA granules down the axon and dendrites (*37*). One possibility is that when secretome mRNA is released from these granules, the presence of lysosomes and Lunapark on a nearby ER junction facilitates local translation of these mRNAs. This might help explain the observed neurodevelopmental defects when Lunapark is dysfunctional (*38, 39*). While we have focused on secretome translation in the cell periphery, lysosomes and Lunapark are likely playing identical roles in secretome translation in the ER of the perinuclear region of cells as well since secretome mRNA translation decreased globally when Lunapark was knocked down. Moreover, we observed Lunapark and lysosomes in high abundance in the perinuclear region of ER, suggesting they are involved in regulating secretome mRNA translation as they do on ER in the cell periphery.

In conclusion, our findings underscore the intricate coordination among ER junctions, Lunapark and lysosomes in regulating the translation of secretome mRNAs. Our proposed model highlights the critical role of Lunapark and lysosomes in organizing subdomains on ER junctions for effective secretome mRNA translation. The arrangement of ER junctions in close proximity to lysosomes enables secretome mRNA translation to be upregulated in response to the release of amino acids and other translation components from lysosomes in an eIF2-dependent manner. Together, our results highlight a novel inter-organellar communication occurring between ER and lysosomes that tunes the level of secretory and membrane protein biosynthesis in cells.

## Acknowledgements

We thank K. Schaefer and D. Walpita for assistance with FACS and cell culture, H. Yi for lentivirus preparation and purification, B. English, C. Gladkova, P. Sengupta, V. Wang, and D. Hwang for insightful discussion, and P. Li and C. Blackstone for the Lunapark-GFP plasmid.

## Author Contributions

H.C., Y.L, Y.J.Y., R.H.S., and J.L-S conceptualized the project. J.G. and L.D.L. synthesized JF dyes, H.C. performed experiments, Y.J.Y. validated L10A-HaloTag function, H.C. and Y.L. designed constructs and experiments for lysosome proximity measurements, H.C. performed computational image analysis and simulations, J.L-S supervised the project. All authors contributed to writing and revising the manuscript and approved the final version.

## Data and materials availability

All materials generated for this study (plasmids, cell lines, etc.) are available upon request from J.L-S. All data and code are also available upon request from J.L-S.

## References

1. D. W. Reid, C. V. Nicchitta, Diversity and selectivity in mRNA translation on the endoplasmic reticulum. Nature Reviews Molecular Cell Biology 16, 221–231 (2015).

2. P. Walter, V. R. Lingappa, Mechanism of Protein Translocation Across the Endoplasmic Reticulum Membrane. Annual Review of Cell Biology 2, 499–516 (1986).

3. R. J. Deshaies, R. Schekman, A yeast mutant defective at an early stage in import of secretory protein precursors into the endoplasmic reticulum. Journal of Cell Biology 105, 633–645 (1987).

4. D. Görlich, E. Hartmann, S. Prehn, T. A. Rapoport, A protein of the endoplasmic reticulum involved early in polypeptide translocation. Nature 357, 47–52 (1992).

5. L. Acosta-Sampson et al., Role for ribosome-associated complex and stress-seventy subfamily B (RAC-Ssb) in integral membrane protein translation. Journal of Biological Chemistry 292, 19610–19627 (2017).

6. L. Bar-Peled et al., A Tumor Suppressor Complex with GAP Activity for the Rag GTPases That Signal Amino Acid Sufficiency to mTORC1. Science 340, 1100–1106 (2013).

7. S. Kim et al., Leucine-sensing mechanism of leucyl-tRNA synthetase 1 for mTORC1 activation. Cell Reports 35, 109031 (2021).

8. S. L. Moon, R. Parker, Analysis of eIF2B bodies and their relationships with stress granules and P-bodies. Scientific Reports 8, (2018).

9. Y. Shibata, G. K. Voeltz, T. A. Rapoport, Rough sheets and smooth tubules. Cell 126, 435–439 (2006).

10. L. Heinrich et al., Whole-cell organelle segmentation in volume electron microscopy. Nature 599, 141–146 (2021).

11. R. S. Lerner, C. V. Nicchitta, mRNA translation is compartmentalized to the endoplasmic reticulum following physiological inhibition of cap-dependent translation. Rna 12, 775–789 (2006).

12. F. M. Fazal et al., Atlas of Subcellular RNA Localization Revealed by APEX-Seq. Cell 178, 473–490.e426 (2019).

13. E. Bertrand et al., Localization of ASH1 mRNA Particles in Living Yeast. Molecular Cell 2, 437–445 (1998).

14. B. Wu, C. Eliscovich, Y. J. Yoon, R. H. Singer, Translation dynamics of single mRNAs in live cells and neurons. Science 352, 1430–1435 (2016).

15. C. Wang, B. Han, R. Zhou, X. Zhuang, Real-time imaging of translation on single mRNA transcripts in live cells. Cell 165, 990–1001 (2016).

16. X. Yan, T. A. Hoek, R. D. Vale, M. E. Tanenbaum, Dynamics of Translation of Single mRNA Molecules In Vivo. Cell 165, 976–989 (2016).

17. T. Morisaki et al., Real-time quantification of single RNA translation dynamics in living cells. Science 352, 1425–1429 (2016).

18. C. J. Obara, A. S. Moore, J. Lippincott-Schwartz, Structural Diversity within the Endoplasmic Reticulum—From the Microscale to the Nanoscale. Cold Spring Harbor Perspectives in Biology 15, a041259 (2023).

19. S. Charrin, S. Jouannet, C. Boucheix, E. Rubinstein, Tetraspanins at a glance. Journal of Cell Science 127, 3641–3648 (2014).

20. S. Chen et al., Lunapark stabilizes nascent three-way junctions in the endoplasmic reticulum. Proceedings of the National Academy of Sciences 112, 418–423 (2015).

21. S. Wang, H. Tukachinsky, F. B. Romano, T. A. Rapoport, Cooperation of the ER-shaping proteins atlastin, lunapark, and reticulons to generate a tubular membrane network. eLife 5, e18605 (2016).

22. K. E. Beatty et al., Fluorescence Visualization of Newly Synthesized Proteins in Mammalian Cells. Angewandte Chemie International Edition 45, 7364–7367 (2006).

23. J.-M. Cioni et al., Late Endosomes Act as mRNA Translation Platforms and Sustain Mitochondria in Axons. Cell 176, 56–72.e15 (2019).

24. Y. Guo et al., Visualizing Intracellular Organelle and Cytoskeletal Interactions at Nanoscale Resolution on Millisecond Timescales. Cell 175, 1430–1442.e1417 (2018).

25. L. Yuniati et al., Ubiquitylation of the ER-Shaping Protein Lunapark via the CRL3KLHL12 Ubiquitin Ligase Complex. Cell Reports 31, 107664 (2020).

26. R. L. Pisoni, J. A. Schneider. (Springer US, 1992), pp. 89–99.

27. U. Bandyopadhyay et al., Leucine retention in lysosomes is regulated by starvation. Proceedings of the National Academy of Sciences 119, e2114912119 (2022).

28. M. Nofal et al., GCN2 adapts protein synthesis to scavenging-dependent growth. Cell Syst. 13, 158–+ (2022).

29. A. G. Hinnebusch, Evidence for translational regulation of the activator of general amino acid control in yeast. Proceedings of the National Academy of Sciences 81, 6442–6446 (1984).

30. M. E. Filbin, J. S. Kieft, Toward a structural understanding of IRES RNA function. Current Opinion in Structural Biology 19, 267–276 (2009).

31. A. Petrov, R. Grosely, J. Chen Seán Joseph, Multiple Parallel Pathways of Translation Initiation on the CrPV IRES. Molecular Cell 62, 92–103 (2016).

32. C. C. Thoreen et al., A unifying model for mTORC1-mediated regulation of mRNA translation. Nature 485, 109–113 (2012).

33. J. C. Tsai et al., Structure of the nucleotide exchange factor eIF2B reveals mechanism of memory-enhancing molecule. Science 359, eaaq0939 (2018).

34. T. Shemesh et al., A model for the generation and interconversion of ER morphologies. Proceedings of the National Academy of Sciences 111, E5243–E5251 (2014).

35. S. Chen, Y. Cui, S. Parashar, P. J. Novick, S. Ferro-Novick, ER-phagy requires Lnp1, a protein that stabilizes rearrangements of the ER network. Proceedings of the National Academy of Sciences 115, E6237–E6244 (2018).

36. D. Akopian, C. A. McGourty, M. Rapé, Co-adaptor driven assembly of a CUL3 E3 ligase complex. Molecular Cell 82, 585–597.e511 (2022).

37. Y.-C. Liao et al., RNA Granules Hitchhike on Lysosomes for Long-Distance Transport, Using Annexin A11 as a Molecular Tether. Cell 179, 147–164.e120 (2019).

38. M. W. Breuss et al., Mutations in LNPK, Encoding the Endoplasmic Reticulum Junction Stabilizer Lunapark, Cause a Recessive Neurodevelopmental Syndrome. The American Journal of Human Genetics 103, 296–304 (2018).

39. L. Ghila, M. Gomez, The evolutionarily conserved gene LNP-1 is required for synaptic vesicle trafficking and synaptic transmission. European Journal of Neuroscience 27, 621–630 (2008).

