## Supplementary Figures and Tables for "Lysosomal release of amino acids at ER three-way junctions regulates transmembrane and secretory protein mRNA translation"

### Materials and Methods

#### *Cell culture and Electroporation*

U-2 OS (ATCC) cells were cultured in phenol-red free DMEM supplemented with 10% fetal bovine serum (Corning), 2 mM L-glutamine (Corning), and 100 IU penicillin and 100 μg/mL streptomycin (CellGro) at 37°C and 5% $CO_2$. For the tet-inducible system, FBS that was tested free of tetracycline (Cytvia) was used. Cells were grown on either diluted fibronectin (1:100, Millipore) or Matrigel (1:100, Corning)-coated coverslip chambers or sterilized cover glasses no. 1.5 Round (EMS). Lonza Electroporation kit with the U-2 OS specific set-up suggested by the company was used for transient transfection, which was performed 16-20 hours prior to imaging. In addition, HeLa (ATCC), COS-7 (ATCC), HEK293T (ATCC), and HT1080 (ATCC) cells were cultured and electroporated according to the manufacturer's instructions.

#### *Plasmids and cloning*

Lists of plasmids used in this study is provided in Supplementary Table 1.

#### *Generation of stable cell line and tet-induction*

To generate lentivirus particles, HEK293T cells were transfected with MCP-GFP, MCP-HaloTag, or scFv-sfGFP plasmids, along with viral packaging plasmids. The supernatant was collected 48-72 hours after transfection, filtered through a 0.22 μm syringe filter, and concentrated using Lenti-X Concentrator (Takara). U-2 OS cells were exposed to lentiviral particles, and MCP-HaloTag and scFv-sfGFP double positive U-2 OS cells were generated via sequential viral transduction and sorted via fluorescence-activated cell sorting (FACS).

To generate cells stably expressing the protein of interest with only MS2 binding sites (MBS) mRNAs, lentiviral transduction was used on pre-sorted MCP-GFP or MCP-HaloTag positive U-2 OS cells. Cells stably expressing a tet-inducible cytERM SUNTAG were generated using the PiggyBac system. MCP-HaloTag/scFv-sfGFP double positive U-2 OS cells were then electroporated with cytERM SUNTAG plasmid with hyperactive piggyBac transposase (hyPBase). After four days post-electroporation, the cells were selected under 10 μg/mL Blasticidin S for two weeks. The stable cells expressing MCP-HaloTag/scFv-sfGFP/cytERM-SUNTAG were induced with 100 ng/mL doxycycline 3-5 hours prior to imaging.

#### *Labeling JF dyes with HaloTag ligand and SNAPTag ligand*

For cells expressing HaloTag, they were incubated in complete media with 100 nM JF646 HaloTag ligand (JF646-HTL) at 37°C for 30 minutes and then washed twice. The rinsed cells were then equilibrated in the media at 37°C for 30 minutes prior to imaging. Similar methods were used for PA-JF646-HTL, PA-JF549-HTL, JF549-HTL, JF646x-HTL, and JF635-HTL.

Cells expressing SNAPTag were incubated in complete media with 250 nM JF549 with chloropyrimidine SNAPTag ligand (JF549-cpSNAP) for 1 hour at 37°C and then washed twice. The rinsed cells were then washed twice again after 30 minutes to remove residual dyes, and then equilibrated in fresh media at 37°C for 30 minutes prior to imaging.

#### *Reagents for Condition Screening*

Rapamycin (Sigma, 2.5 mg/mL in DMSO), Torin-1 (Cell Signaling, 1 mM in DMSO), Cycloheximide (Sigma, 100 mg/mL in DMSO), Chloroquine (Sigma, 10 mM in $H_2O$),

Thapsigargin (Sigma, 1 mM in DMSO), Protease cocktail inhibitor (ThermoFisher) and Puromycin (ThermoFisher, 10 mg/mL in $H_2O$). Amino acid-free medium is composed of Amino acid free DMEM (USBiological), 10% dialyzed FBS (ThermoFisher), 4.5 g/L Glucose (Sigma), 110 mg/L Sodium Pyruvate (Sigma), 3.7 g/L Sodium Bicarbonate (Sigma), 2 mM L-Glutamine (Corning), 100 IU penicillin and 100 µg/mL streptomycin (CellGro).

*Single Molecule Tracking with HILO Microscopy*

Single molecule tracking of single ribosomes was conducted using a customized Nikon TiE inverted microscope with a split port for simultaneous imaging onto two Andor iXON DU-897 EM-CCD cameras, equipped with a 100x Apo TIRF 1.49 oil-immersion objective (Nikon). U-2 OS cells were co-transfected with L10A-HaloTag plasmid and either mEmerald Sec61b or SNAPf-Sec61b plasmids and imaged after 16 hours. These cells were labeled with either PA-JF549-HTL or PA-JF646-HTL/JF549-cpSNAP (*1*). A brief pulse of 405 nm laser was used to excite and activate the PA-JF dyes. The incident angle for excitation lasers was manipulated to achieve HILO illumination. The emission resulting from simultaneous excitation of two fluorophores was split using either T647LPXR (Chroma) or T565LPXR (Chroma) and collected in two Andor iXon EM-CCDs with emission filters: ET525/50m (Chroma) for GFP, ET605/70m (Chroma) for PA-JF549 or JF549, and ET700/75m (Chroma) for PA-JF646.

*Single Particle Tracking (SPT) with Spinning Disk Confocal*

Single molecule tracking for single mRNAs was conducted using a customized Nikon TiE inverted microscope, outfitted with a Yokogawa spinning-disk scan head (#CSU-X1, Yokogawa) and Andor iXON EM-CCD cameras. Fluorescence was collected using a 100x Apo TIRF 1.49 oil-immersion objective (Nikon). For live-cell imaging, cells were imaged in complete medium and incubated with a stage heater (Tokai Hit) at 37°C and 5% $CO_2$. Two-color imaging was performed similarly to single molecule tracking, as described above. Three-color imaging was performed using trigger mode. The emission from 561 and 640 nm excitation was collected sequentially into the same camera using a multi-bandpass emission filter (Chroma, ZET405/488/561/647m).

*Immunofluorescence and Proximity ligation Assay*

For immunofluorescence labeling, U-2 OS cells stably expressing MCP-HaloTag/scFv-sfGFP/cytERM-SUNTAG were plated on matrigel-coated 12 mm coverslips (EMS) in sterile 24-well tissue-culture graded plates (Corning). The cells were induced with 100 ng/mL doxycycline (Sigma) and labeled with JF dyes as described above. The cells were then fixed with 4% (w/v) paraformaldehyde (PFA) + 0.2% glutaraldehyde for 15 minutes at room temperature (RT). Cells were permeabilized with 0.1% Triton X-100 for 10 minutes, then treated with 1% BSA (Sigma) in phosphate-buffered saline (PBS) plus 0.05% Tween-20 (PBS-T) for 1 hour. The primary antibody (Rabbit Anti-LNPK antibody, Sigma) was added overnight at 4°C and washed with 1% BSA + PBS-T. The coverslip was incubated with secondary antibodies conjugated with Alexa Fluor 594 (ThermoFisher) for 1 hour at RT and rinsed with PBS. The coverslip was mounted on a slide and imaged using a Zeiss 980 AiryScan equipped with a 63x objective.

Proximity Ligation Assay (PLA) was performed using the Duolink In situ Orange Starter kit Mouse/Rabbit (Sigma). U-2 OS cells were plated on Matrigel-coated 12 mm coverslips (EMS) in sterile 24-well tissue-culture graded plates (Corning). The manufacturer's protocol was followed, using the following antibodies: Rabbit LNPK antibody (Sigma, HPA014205-25), Mouse EEA1 antibody (BD Biosciences, 610456), Mouse LAMP1 antibody (Abcam, ab25630), Mouse TOMM20 antibody (66777-1-Ig).

*Immunoblotting*

For immunoblotting, U-2 OS cells were collected via scraping the plate in lysis buffer (25 mM Tris-HCl pH 7.6, 150 mM NaCl, 1% NP-40, 1% sodium deoxycholate, 0.1% SDS) supplemented with EDTA-free protease inhibitor cocktail (Roche) and PhoSTOP (Roche), followed by sonication. The lysate was quantified with a BCA kit (ThermoFisher), following the manufacturer's protocol and diluted using 4X NuPage LDS Sample buffer (ThermoFisher) supplemented with 0.1 M DTT (Sigma Aldrich). The samples were heated in a 95°C heat block for 10 minutes. Equal amounts of lysate (~ 5 µg) were run on 4-12% NuPage Bis-Tris Mini protein gels (ThermoFisher). After electrophoresis, the proteins were transferred onto 0.2 µm Nitrocellulose membranes using the Trans-Blot Turbo system. The membranes were blocked using EveryBlot blocking buffer (Bio-Rad). Primary antibodies were diluted 1:1000 in the same blocking buffer and incubated overnight at 4°C with gentle agitation. After washing three times with TBS-T, secondary antibody staining was performed for 1 hour at room temperature. The membranes were washed with TBS-T and developed with SuperSignal West Pico PLUS Chemiluminescent Substrate (ThermoFisher). The membranes were imaged on ChemiDoc (Bio-Rad) with optimized exposure for signal detection.

*Hybridization chain reaction single molecule fluorescence in situ hybridization (HCR smFISH)*

For HCR smFISH, we designed and purchased oligonucleotides that hybridize to endogenous Human CD9 mRNA from Molecular Instruments. U-2 OS cells transiently transfected with mEmerald-Sec61b were plated on Matrigel-coated 12 mm coverslips in sterile 24-well tissue-culture graded plates. Cells were fixed with 4% (w/v) PFA + 0.2% glutaraldehyde for 15 minutes at RT. Hybridization was performed based on the manufacturer's protocol for mammalian cells on a coverslip for single molecule detection. The coverslips were mounted on a glass slide and imaged immediately.

*Single Particle Tracking Analysis*

L10A-HaloTag and mRNA tracking was performed using TrackMate software (FIJI). The resulting trajectories were imported into Matlab using a custom Matlab code to store x, y, and t. Displacement of each trajectory was calculated, and each trajectory was categorized based on the mask to classify cytosolic trajectories and remove nuclear localizations using custom Matlab code.

The mean squared displacement (MSD) of the population was calculated by averaging squared displacements (SDs) at a given lag time. The apparent diffusion coefficient ($D_{app}$) was calculated based on a linear fit of MSD vs lag time using the built-in linear fit module in Matlab, described by Equation 1.

$$MSD(\tau) = 2nD_{app}\tau + 4\sigma^2 \quad \text{(eq.1)}$$

where $D_{app}$ = diffusion coefficient, $\tau$ is lag time, n = number of dimensions (2), $\sigma$ = localization error. The first four points were used to fit the curve to estimate the apparent diffusion coefficient as many of these trajectories showed a confined diffusion.

Mean square displacement from each trajectory ($MSD_T$) was calculated by averaging the squared displacement (SD) within the same trajectories. For determining the translationally active population, we analyzed trajectories that are longer than 9-step at 1 Hz and averaged 9 individual SDs to generate $MSD_T$, as described in Equation 2.

$$MSD_{\tau} = \frac{\sum_{i=1}^{n}((x_{i+1}-x_i)^2+(y_{i+1}-y_i)^2)}{n} \quad \text{(eq.2)}$$

*Monte Carlo Simulation of Brownian Motion*

Monte Carlo simulation of random walk was performed using a custom-built Matlab code. For each time step, individual particles choose displacement in each of the three Cartesian directions, and its distribution is defined by the free, 3D diffusion coefficient D. The squared displacement of the corresponding Gaussian propagator is sqrt(2Dt).

*Motion-based analysis of translating mRNA*

Displacements of single trajectories of mRNAs in the cytosol are calculated based on their time progression. Each trajectory longer than 10-step was used to calculate the mean square displacement in 1 second. Each trajectory was then categorized based on the MSD cut-off of 0.055 $\mu m^2$ based on Equation 2. For determining the translation fraction, we calculated by dividing the number of trajectories that are classified as translation by the number of total classified trajectories.

To generate pseudo-colored single molecule movies, the positions and time-mark from categorized trajectory was mapped onto two blank matrices (x,y,t) where one represents translationally active and the other represents translationally silent mRNAs. The resulted single pixel movies were saved as tagged image file (tif). The gaussian blurring of 50 nm was applied to mimic the localization accuracy of each molecule. The resulted movie was then overlayed on the images of corresponding ER (Sec61b).

*SUNTAG intensity analysis and FRAP*

Foci of $scFv_{GCN4}$-sfGFP signal were localized and quantified using TrackMate. The background signal was subtracted and each (x,y,t) coordinate was correlated to the trajectories of corresponding mRNAs so that the maximum distance away is less than or equal to 2 pixels. Then, only mRNAs that are in the cytoplasm were selected for analysis. We calculated the translating fraction of mRNA by dividing SUNTAG(+) mRNAs over all tracked mRNAs in the cytoplasm.

Fluorescence recovery after photobleaching (FRAP) experiments were perform on a customized Nikon TiE inverted scope equipped with a Yokogawa spinning-disk scan head (#CSU-X1, Yokogawa) and Andor iXON EMCCD cameras. The excitation and emission was performed on 100x Apo TIRF 1.49NA oil-immersion objective (Nikon). Photobleaching was performed using a Bruker Mini-scanner that scans the region of interest. Sequential excitation of 488 and 640 nm laser was performed to collect scFv-sfGFP and MCP-HaloTag signal respectively through a multi-bandpass emission filter (Chroma, **ZET405/488/561/647m)**. The cells were imaged prior to bleaching for 10 consecutive frames and imaged post-bleaching at 5 or 10 seconds per frame for >5 minutes. Particles were tracked using Imaris spot tracker (Oxford Instrument) and corresponding bleached SUNTAG signal was analyzed over time.

*Lysosome-SUNTAG correlation*

Lysosomes were tracked using TrackMate with the proper threshold to detect the center of the lysosome. The distance from the center of lysosome to each mRNA positions were calculated. The minimum distance from lysosome to each mRNA was then used for analysis. SUNTAG signals that were quantified during the SUNTAG intensity analysis was referenced to the minimum lysosome distance. The histogram of average SUNTAG intensity vs average minimum lysosome distance was quantified by binning the lysosome distance at 500 nm interval, starting from 750 nm to 4.25 µm. The SUNTAG intensity of the corresponding average lysosome distance binned at a 500-nm interval starting from 750 nm to 4.25 µm was used to generate the histogram.

*siRNA knockdown and KD efficiency*

25 pmol of siRNAs targeting Lunapark or CLIMP63 (Dharmacon) were transfected into U-2 OS cells using Lipofectamine RNAiMAX (ThermoFisher) in Opti-MEM (ThermoFisher) in a 6-well chamber (Corning). The transfected cells were incubated for 48 to 72 hours before imaging. Prior to imaging, the total RNA of the cells from same-day transfection was collected using RNA extraction kits (NEB). The concentration of total RNA was measured using UV spectrometer (ThermoFisher) and performed One-Step RT-qPCR (NEB) on Roche LightCycler. Knockdown efficiency was calculated based on calculating $\Delta\Delta C_q$ using Actin mRNA as a standard. The efficiency was plotted in Fig. S7 The list of primers used for this is listed in Supplementary Table 2.

*CRISPR Knockout of Lunapark*

To generate cells with targeted knockout of human Lunapark, single guide RNAs (sgRNAs) were designed against human Lunapark from Synthego. The sgRNA sequences (AGCAAAAAAUGGGAGUGUCA, UGUAAACAGAUAGAGAACUG, UCAAGCAUUGGAAGAAUUUA) were then assembled with SpCas9 2NLS Nuclease (Synthego) as per the manufacturer's instructions. The resulting sgRNA-Cas9 complex was then electroporated into U-2 OS cells using a Lonza 4D nucleofector programmed for U-2 OS cells. The cells were expanded and cultured for single colonies by diluting them in 96-well plates (Corning). The genome of each expanded colony was then collected using the Monarch Genomic DNA purification kit, and PCR was performed on the purified DNA using primers provided by Synthego (listed in the Supplementary Table 3). The resulting Sanger sequencing data was analyzed using Synthego software to identify positive colonies. These colonies were then checked for Lunapark expression by collecting the lysate and performing immunofluorescence or immunoblotting as described above.

*HPG Incorporation Assay*

The Click-It HPG Alexa Fluor 488 protein synthesis kit (ThermoFisher) was used to perform the HPG incorporation assay. Prior to the addition of HPG (50 µM), U-2 OS cells were incubated with methionine-free DMEM for 1 hour. HPG was incorporated into the cells over a period of 1.5 hours. As HPG can also be incorporated into mitochondria, a cycloheximide incubation control was incorporated for all tested conditions. To monitor HPG incorporation into the membrane fraction only, HPG-incorporated cells were incubated with a solution containing 0.025% Digitonin, 115 mM KAc, 25 mM HEPES pH 7.4, 2.5 mM $MgCl_2$, 2 mM EGTA, and 150 mM sucrose for 3 minutes at 37°C and proceeded with fixation. For the whole cell fraction, the cells were fixed using 4% PFA and 0.2% glutaraldehyde for 15 minutes, and the washing and labeling protocol followed the manufacturer's instructions. The labeled cells were mounted using VECTASHIELD antifade with DAPI (Vectorlabs), imaged using Zeiss 980 AiryScan, and quantified using CellProfiler. To remove the effect from mitochondrial translation, the quantified signal of each condition was subtracted with the corresponding cycloheximide treatment.

**Supplementary Table 1.** List of Plasmids for this study

| **Name** | **Source** |
|---|---|
| pUBC-SiT-EGFP-MS2 | This study |
| pUBC-Calreticulin-mEmerald-MS2 | This study |
| pUBC-CD4-EGFP-MS2 | This study |
| pUBC-Halo-Actin-MS2 | (*2*) |
| pCMV-SNAPf-Sec61b | This study |
| pCMV-mApple-Sec61b | (*3*) |
| pUBC-L10A-Halo | This study |
| pPGK-Lamp1-mApple | (*4*) |
| pPGK-Lamp1-mScarlet-i | This study |
| pPB-cytERM-SUNTAG-MS2 | This study |
| pPB-CrPV-IRES-cytERM-SUNTAG-MS2 | This study |
| pUBC-MCP-Halo | (*2*) |
| pUBC-MCP-GFP | (*2*) |

**Supplementary Table 2**. siRNA and qPCR primers used for this study

| **siRNA sequences used in this study** | |
|---|---|
| *CLIMP63 siRNA SMARTpool Sequences* | |
| siRNA CKAP4, D-012755-01 | CCAAAUCCAUCAACGACAA |
| siRNA CKAP4, D-012755-02 | CCAAGGGUUUACUAGAUGA |
| siRNA CKAP4, D-012755-03 | AGACCCAGCUGGUGCUCUA |
| siRNA CKAP4, D-012755-04 | GAGCGUGCAUACUGCGUUU |
| *Lunapark siRNA SMARTpool Sequences* | |
| siRNA KIAA1715, D-023148-01 | GGUAUGCACUUAUAUGUCA |
| siRNA KIAA1715, D-023148-02 | AUUAUGGGUUGGAAGAUUA |
| siRNA KIAA1715, D-023148-03 | ACGAUGUUCUUGAUGAUAA |
| siRNA KIAA1715, D-023148-04 | CGGCUAAAUUAAUUCUUGA |
| qPCR Primer sets | |
| *Lunapark (LNPK)* | |
| Forward Set 1 | CTC CTC CAC AAG TTC CAG TAT |
| Reverse Set 1 | GAT GAT AGG GCT GGA GTA ACA G |
| Forward Set 2 | ACT GTT ACT CCA GCC CTA TCA |
| Reverse Set 2 | CTA TCC AAA GCA CCT CGT TCT C |
| Forward Set 3 | GCA AGA CCT GGA CAA GAG ATT |
| Reverse Set 3 | TTG GTG GTC CAG GAG ATA CT |
| *CLIMP63 (CKAP4)* | |
| Forward Set 1 | GGA ATC AGC CAA GGG TTT ACT A |
| Reverse Set 1 | CCT GCA CAC GCA ATT CAT TTA |
| Forward Set 2 | GGA AGC TGT GAA GGA GAT ACA G |
| Reverse Set 2 | ACC TCG GTG TAG ATG TCA GA |
| Forward Set 3 | TGG TTG CAT ACT CGG TCA AA |
| Reverse Set 3 | CCA GAT CAT TCC TCA GGT CAT C |
| *β-Actin (ActB)* | |
| Forward Set 1 | CAC CAT TGG CAA TGA GCG GTT C |
| Reverse Set 1 | AGG TCT TTG CGG ATG TCC ACG T |

**Supplementary Table 3**. Single guide RNAs for generating Lunapark KO U-2 OS cells

| **Lunapark KO Generation** | |
|---|---|
| *gRNA sequence targeting Human Lunapark* | |
| sgRNA-LNPK-1 | AGCAAAAAAUGGGAGUGUCA |
| sgRNA-LNPK-1 | UGUAAACAGAUAGAGAACUG |
| sgRNA-LNPK-1 | UCAAGCAUUGGAAGAAUUUA |
| *Amplification and sequencing primers* | |
| KIAA1715 Forward-1 | CCTCAGGTATATTGGATTACAAATGAAGG |
| KIAA1715 Reverse-1 | TGCCGATACCAATTTCATTCCC |
| KIAA1715 Seq-1 | CCTCAGGTATATTGGATTACAAATGAAGG |
| KIAA1715 Reverse-2 | TCATTCCCTGAATCTTTGCCA |

**Supplementary Methods**

The probability distribution of effective 2D diffusion coefficient, $p(D_{eff})dD_{eff}$ where $D_{eff}$ = $<r^2>/(4*i\Delta t)$, $<r^2>$ = mean squared displacement, and $i\Delta t$ = time lag, is described in (*5*) and with Equation 1 (Eq.1)

$$p(D_{eff}) = \frac{1}{(N-1)!}(\frac{N}{D_0})^N (D_{eff})^{N-1} e^{-\frac{N D_{eff}}{D0}} \quad \text{(Eq.1)}$$

where N = $N_{total}/ i\Delta t$, $N_{total}$ = length of trajectory, $D_0$ = true diffusion coefficient, $D_{eff}$ = effective diffusion coefficient for an individual trajectory based on its squared displacement. As N approaches, the distribution of $D_{eff}$ becomes tighter. Assuming that translation is highly processive and that there is no state changing within the time from of our imaging, translationally active and silent mRNAs can be described as two independent populations whose diffusive motions can be described with two $D_0$'s.

We performed Monte Carlo simulation to understand the required separation for N-averaging based on the free 2-D diffusion. The Monte-Carlo simulations of 5000 trajectories with two different diffusion coefficients, 0.007 $\mu m^2 s^{-1}$ (Blue) and 0.033 $\mu m^2 s^{-1}$ (Orange) are performed. These diffusion coefficients are based on the $D_{app}$ of SUNTAG(+) mRNA and $D_{app}$ of puromycin-treated SiT-EGFP mRNA, respectively. The square displacement from lag time of 1 second was calculated for each trajectory. Then, N number of steps that are averaged in each trajectory and populated into a separate histogram plot (**Fig. S2**). The false positive (FP) rate is calculated based on how many trajectories based on N-step averaging belongs to the set cut-off of 0.055 $\mu m^2$ (red dotted line).

A typical MSD vs lag time distribution for both SUNTAG+ mRNA and SiT-EGFP mRNA shows a confined diffusion, as the initial slope overestimates MSD with increased lag time. Therefore, the comparison needs to be done at the same time scale to avoid the effect. This method most likely overestimates the true translating fraction of mRNA as the longer than 9-step trajectories are required for this analysis to perform properly whereas the average trajectory length of MS2-labeled translocating protein mRNA is ~7-8 steps. Translationally silent (TS) mRNAs presumably will have a shorter length of trajectories. Hence, these molecules are likely be missed during tracking. Fortunately, we observed that puromycin-treatment at the same time scale appears to predict significantly reduced population of translationally active SiT-EGFP mRNAs, which suggests that this method can detect the changes in translation fraction. Since motion-based analyses are prone to other artifacts, such as increased directional motions or structural changes to ER, we tried to compare most of our results with quantifications of SUNTAG+ cytERM-23xGCN4 mRNAs.

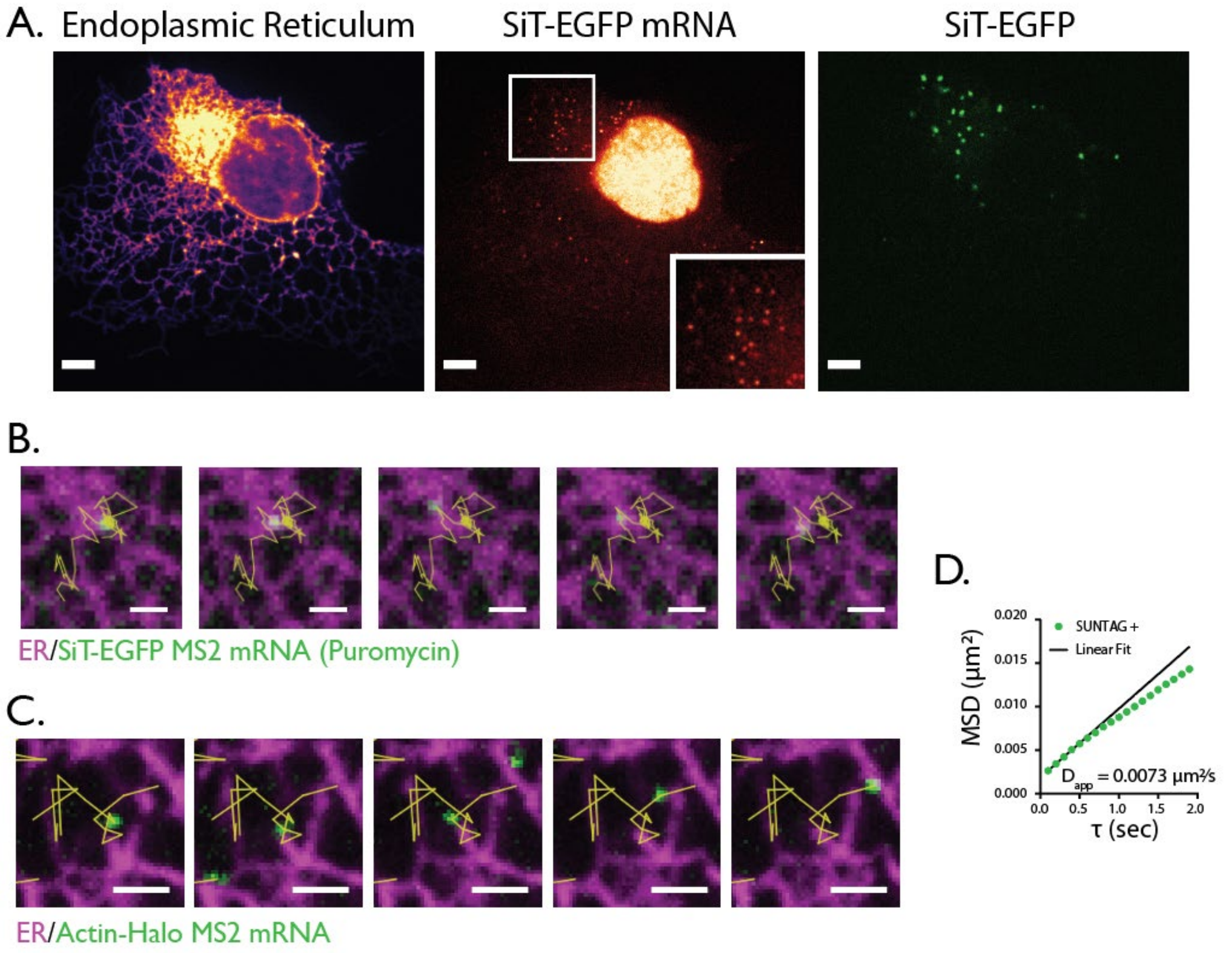

**S1. A**) Golgi-like distribution of SiT-EGFP MS2 mRNA. SiT-EGFP (Green), MS2 (Magenta). The scale bar shows 3 µm. **B)** Time-lapse images of Puromycin-treated SiT-EGFP mRNA. The scale bar represents 1 µm. **C)** Time-lapse images of Actin-Halo mRNA. The scale bar represents µm **D)** Mean Squared Displacement (MSD) vs lag time for SUNTAG(+) cytERM-SUNTAG MS2 mRNA. The linear fit shows $D_{app}$= 0.007 µm$^2$/s

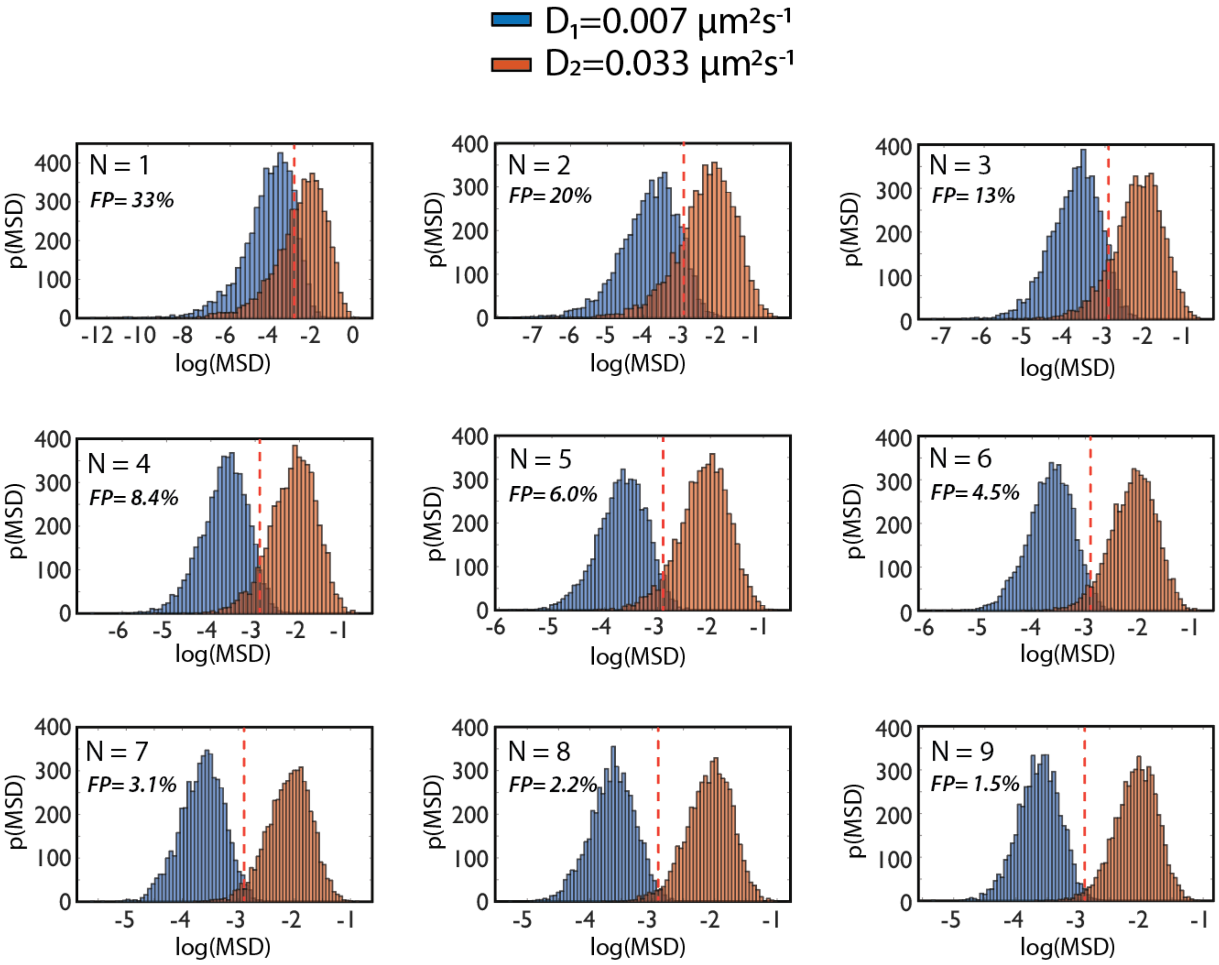

**S2**. Probability distribution of mean squared displacement (MSD) at different N-step averaging of simulated trajectories with $D_1$ (blue, 0.007 µm$^2$s$^{-1}$) and $D_2$ (orange, 0.033 µm$^2$s$^{-1}$). False positive (FP) rates were determined based on the cut-off at 0.055 µm$^2$ (red dotted line).

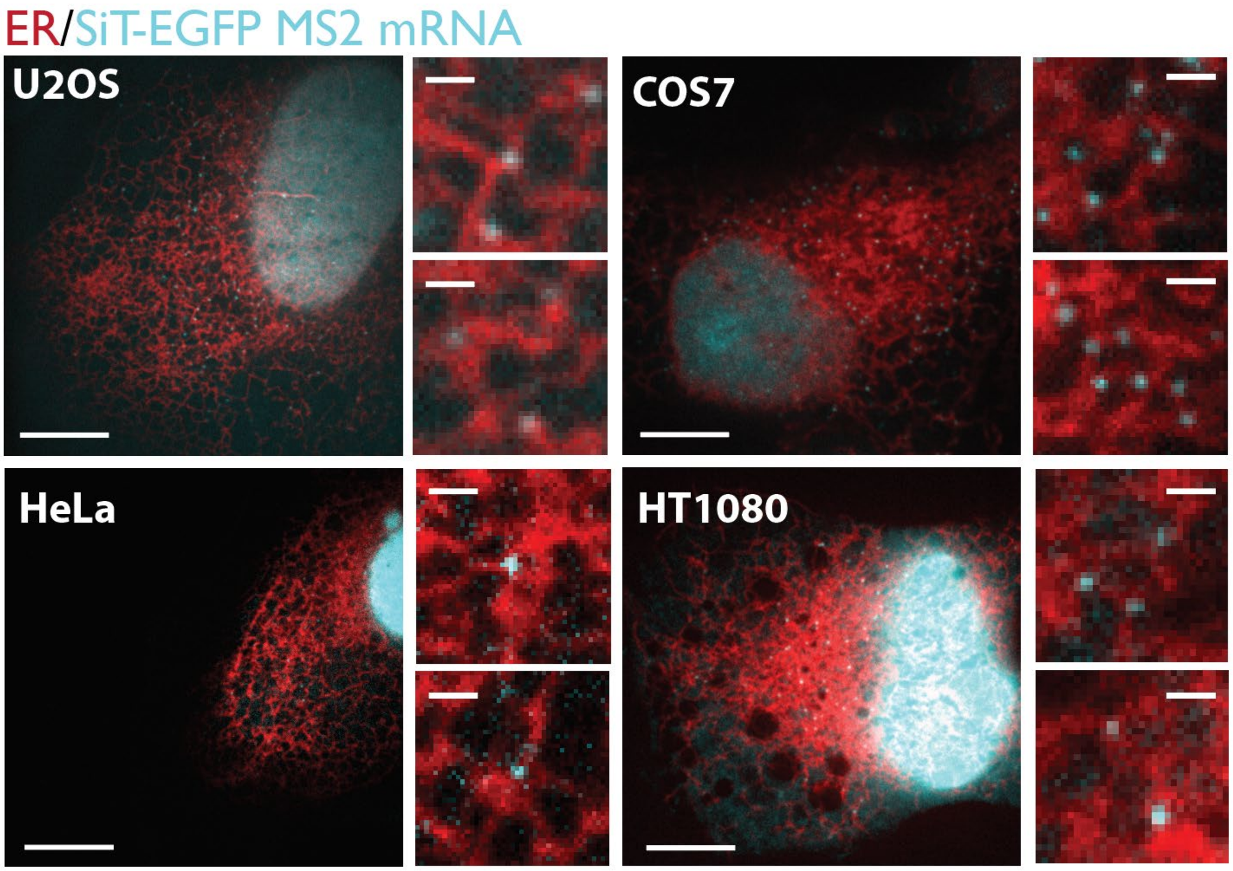

**S3**. Images of U2OS, COS7, HeLa, and HT1080 with SiT-EGFP MS2 mRNA (cyan) and ER (red). The scale bar of large images represents 10 μm. The scale bar of the inset represents 1 μm.

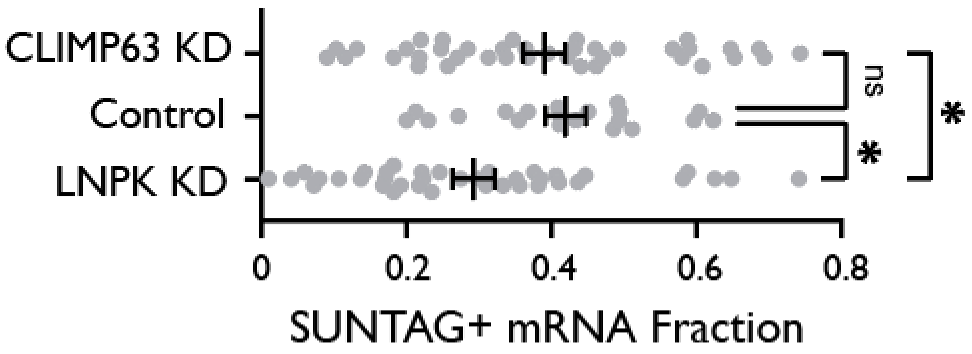

**S4.** SUNTAG(+) cytERM-SUNTAG MS2 fraction in CLIMP63 KD, LNPK KD and control U-2 OS cells. The statistical comparison was perform against control condition.

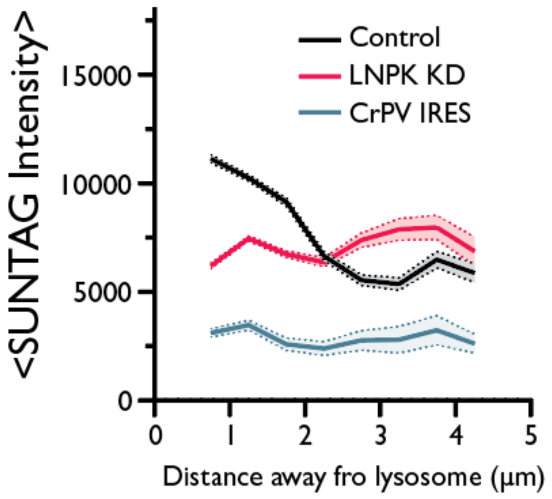

**S5.** Averaged absolute SUNTAG intensity of cytERM-SUNTAG MS2 mRNA over the distance away from the lysosome in µm.

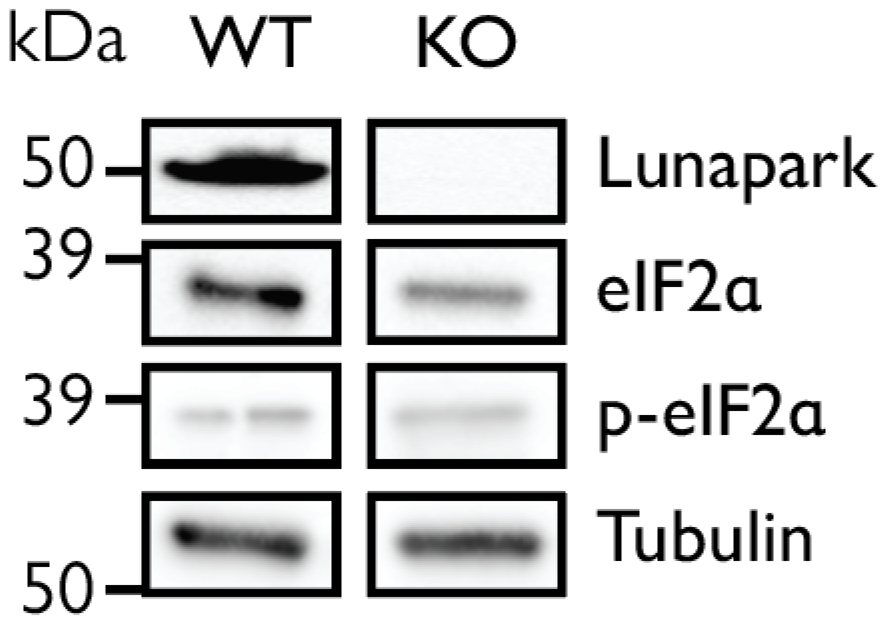

**S6.** Western blot images probing for Lunapark, eIF2α, phosphorylated-eIF2α, and Tubulin in parallel blots using same imaging conditions.

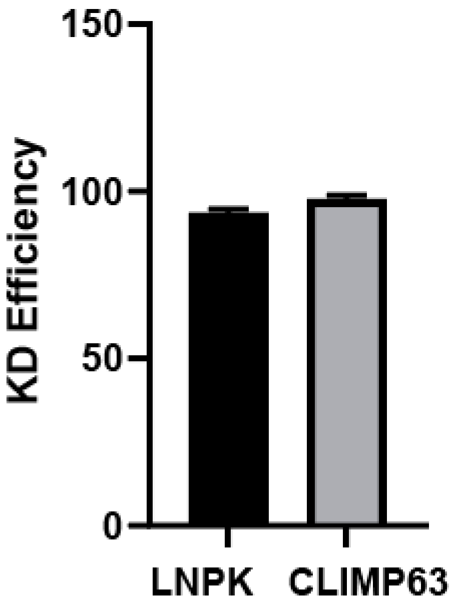

**S7.** siRNA knockdown efficiency in U-2 OS cells measured by qRT-PCR by $\Delta\Delta C_q$ against beta-actin mRNA as a standard.

**Supplementary Movie 1**
SiT-EGFP MS2 mRNA (red) on ER marked by SNAPf-SEC61B (cyan) in U-2 OS cells. Time is in second. The scale bar represents 5 μm.

**Supplementary Movie 2**
Halo-Actin MS2 mRNA (red) on ER marked by SNAPf-SEC61B (cyan) in U-2 OS cells. Time is in second. The scale bar represents 5 μm.

**Supplementary Movie 3**
SUNTAG(+) cytERM-SUNTAG MS2 mRNA movie. Anti-GCN4 scFv-sfGFP (SUNTAG, Green) and MCP-Halo (RNA, Magenta). Time is in second. The scale bar represents 1 μm.

**Supplementary Movie 4**
SUNTAG(-) cytERM-SUNTAG MS2 mRNA movie. Anti-GCN4 scFv-sfGFP (SUNTAG, Green) and MCP-Halo (RNA, Magenta). Time is in second. The scale bar represents 1 μm.

**Supplementary Movie 5**
Time-lapse imaging of ISR inhibitor (ISRIB) treatment on Thapsigargin-treated U-2 OS cells expressing cytERM-SUNTAG MS2 mRNA. 200 nM ISRIB was added at t = 0 min. SUNTAG signal (Cyan) and mRNA (Magenta). Time is in minute. The scale bar represents 10 μm.

**Supplementary Movie 6**
Time-lapse imaging of ISR inhibitor (ISRIB) treatment on amino acid deprived U-2 OS cells expressing cytERM-SUNTAG MS2 mRNA. 200 nM ISRIB was added at t = 0 min. SUNTAG signal (Cyan) and mRNA (Magenta). Time is in minute. The scale bar represents 10 μm.

### References:

40. J. B. Grimm *et al.*, Bright photoactivatable fluorophores for single-molecule imaging. *Nature Methods* **13**, 985-988 (2016).
41. Y. J. Yoon *et al.*, Glutamate-induced RNA localization and translation in neurons. *Proceedings of the National Academy of Sciences* **113**, E6877-E6886 (2016).
42. J. Nixon-Abell *et al.*, Increased spatiotemporal resolution reveals highly dynamic dense tubular matrices in the peripheral ER. *Science* **354**,  (2016).
43. M. Vrljic, S. Y. Nishimura, S. Brasselet, W. E. Moerner, H. M. McConnell, Translational Diffusion of Individual Class II MHC Membrane Proteinsin Cells. *Biophysical Journal* **83**, 2681-2692 (2002).
